# Zebrafish Kit ligands cooperate with erythropoietin to promote erythroid cell expansion

**DOI:** 10.1101/2020.02.03.931634

**Authors:** Jana Oltova, Ondrej Svoboda, Olga Machonova, Petra Svatonova, David Traver, Michal Kolar, Petr Bartunek

**Author notes:** Corresponding author: Petr Bartunek, Institute of Molecular Genetics of the Czech Academy of Sciences Videnska 1083, 142 20 Prague 4, Czech Republic.

## Abstract

Kit ligand (Kitlg) is pleiotropic cytokine with a prominent role in vertebrate erythropoiesis. Although the role of Kitlg in this process has not yet been reported in *Danio rerio* (zebrafish), in the present study, we show that its function is evolutionary conserved. Zebrafish possess two copies of Kitlg genes (Kitlga and Kitlgb) due to whole genome duplication. To determine the role of each ligand in zebrafish, we performed a series of *ex vivo* and *in vivo* gain- and loss-of-function experiments. First, we tested the biological activity of recombinant Kitlg proteins in suspension culture from zebrafish whole kidney marrow and we demonstrate that Kitlga is necessary for expansion of erythroid progenitors *ex vivo*. To further address the role of *kitlga* and *kitlgb* in hematopoietic development *in vivo*, we performed gain-of-function experiments in zebrafish embryos, showing that both ligands cooperate with erythropoietin (Epo) to promote erythroid cell expansion. Finally, using the *kita* mutant (*kita*^*b5*/*b5*^ or *sparse*), we show that Kita receptor is crucial for Kitlga/b cooperation with Epo in erythroid cells. In summary, using optimized suspension culture conditions with recombinant cytokines (Epo, Kitlga), we are reporting for the first time *ex vivo* suspension cultures of zebrafish hematopoietic progenitor cells, which can serve as an indispensable tool to study normal and aberrant hematopoiesis in zebrafish. Furthermore, we conclude that although partial functional diversification of Kit ligands has been described in other processes, in erythroid development, both paralogs play a similar role and their function is evolutionary conserved.

****Key points:**:**

- Kit signaling contributes to erythroid cell development and is conserved from fish to man
- Ex vivo expansion and self-renewal of zebrafish erythroid progenitors requires addition of recombinant Kitlga

## Introduction

The most prominent cytokines that regulate proliferation and differentiation of erythroid cells in vertebrates are erythropoietin (EPO)^1–3^ and Kit ligand (KITLG or stem cell factor, SCF)^4–9^. As was described in other vertebrates but not in zebrafish, binding of EPO and KITLG to their cognate receptors ensures erythroid lineage commitment by triggering specific signaling events^1,8,10,11^.

Besides its role in erythropoiesis, KITLG acts pleiotropically, affecting a wide range of tissues and cells, including hematopoietic (HSCs) and germ stem cells ^12–14^. It is an important regulator that plays a role in many processes, both during ontogenesis and in the adult organism^13,15^. In mammals, KIT signaling is associated with erythro-myelopoiesis^16^, as well as neurogenesis and pigmentation^13,17–19^. Interestingly, there are two forms of KITLG occurring *in vivo* – transmembrane, important for the regulation of stem cells in their niches, and soluble, affecting more distant tissues^13,20^. Binding of KITLG to its receptor KIT, a member of receptor tyrosine kinase type-III family, leads to its autophosphorylation, further triggering various signaling cascades, including PI-3K, MAPK, SRC and JAK kinase pathways^20,21^.

In teleost species that include zebrafish, this scenario is more complex due to the fact that the whole ligand-receptor signalosome has been duplicated as a result of an extra round of whole genome duplication. Future studies are therefore dependent on the understanding of diversification of the functions of both ligands (Kitlga, Kitlgb) and receptors (Kita, Kitb) and their binding specificities. Similarly, there are also two copies of the *epo* gene present in the zebrafish genome - *epoa* and *epob*. The zebrafish *epoa* seems to play a similar role as its mammalian ortholog^22,23^ but no role in hematopoiesis has been so far reported for *epob (OS, PB, unpublished).* For simplicity, we will use the designation Epo/*epo* for Epoa/*epoa* in this study.

Current data support the hypothesis that Kit receptor and Kit ligand paralogs have subspecialized during evolution. Kita is expressed in the neural crest, lateral line and notochord^24^. Overexpression of Kitlga results in hyper-pigmentation^25^ and *kita* receptor mutants (*kita*^*b5*/*b5*^ or *sparse*) have defective pigmentation^24^. On the other hand, the second zebrafish Kit paralog, Kitb, is expressed in neural tube and otic vesicles and likely does not play a role in melanogenesis^26,27^. Neither has a role in melanogenesis been reported for Kitlgb^27^.

Although studied extensively, the role of Kit signaling in zebrafish hematopoiesis has remained unknown for a long time. So far, there are only two studies that suggest potential roles of Kitlg in hematopoiesis in *D. rerio*. The first shows a mild increase of HSCs upon overexpression of Kitlgb^28^, the second reports a decrease in the number of HSCs upon downregulation of Kitb; however, no such phenotype was observed for Kita or Kitlga. Based on that, the authors concluded that neither Kita nor Kitlga are involved in hematopoiesis, but only in melanocyte formation, as suggested by previous studies^27^. Contradictory to the findings in other vertebrate models, and despite the fact that Kit receptors are expressed in hematopoietic tissues^24,27^, hematopoiesis is not affected in the adult *kita* mutants (*kita*^*b5*/*b5*^ or *sparse*)^24^ under steady-state conditions.

The utilization of zebrafish as a model organism requires understanding of the regulatory mechanisms of hematopoietic lineage development and their evolutionary conservation, as well as proper approaches to study blood development and disease. The objective of this study is to better understand the importance of Kit ligands in zebrafish hematopoiesis, and thus providing important insights for further research of hematopoietic development. Although *ex vivo* expansion of erythroid progenitors has been reported in different vertebrate organisms^29^, a similar approach that would enable analysis of normal and aberrant hematopoiesis in zebrafish is not yet available. Here, we establish suspension culture conditions for the expansion of zebrafish erythroid progenitors and we investigate the role of the two zebrafish paralogs of Kit ligands (Kitlg) in zebrafish hematopoiesis using *in vivo* and *ex vivo* experimental approaches.

## Methods

### Animal stocks and embryos

Fish were mated, raised and staged according to “The zebrafish book. A guide for the laboratory use of zebrafish (*Danio rerio*)”^30^ and the recommendations for zebrafish husbandry and housing^31^. For tracking the animals (age, number, health, genotype), we used an in-house developed dedicated database solution called Zebrabase^32^. Fish were kept in ZebTEC aquatic system (Tecniplast). Transgenic reporter lines expressing fluorescent genes under the control of tissue specific promoters (*gata1*:*DsRed*^33^ and *lcr:EGFP*^34^, as well as mutant lines (kita^b5/b5^ or *sparse*)^24^ and wild-type animals WT(AB) were used in this study. For *ex vivo* experiments, 6-month old fish were used to ensure optimal number and state of whole kidney marrow cells. Animal care and experiments were approved by the Animal Care Committee of the Institute of Molecular Genetics, Czech Academy of Sciences (13/2016 and 96/2018) in compliance with national and institutional guidelines.

### Ex vivo cultures

Zebrafish whole kidney marrow cells were isolated as previously described^35^. Cells were fractionated using Biocoll (1.077 g/ml, Merck L6115) density centrifugation and initially seeded at 3×10^6^ cells/ml and cultivated in zfS13 medium (for composition, see Supplemental Table 3) at 32°C and 5% CO_2_. Carp serum was obtained by the procedure described and published previously^35^. Specific cytokines (Epo, Gcsfa, Kitlga and Kitlgb) were added at final concentrations of (100 ng/ml), dexamethasone (Dex) was added at final 1μM. Cells were counted using CASY Cell Counter & Analyzer (OMNI Life Science). Following days, cells were maintained at 2×10^6^ cells/ml and one third of the medium containing fresh cytokines and Dex was exchanged every other day to ensure optimal growth of the cells. To study the clonogenic potential of the whole kidney marrow cells after addition of different cytokines, we performed clonal assays in semisolid media (methylcellulose), which prevents movement of the single cells plated in it. For the detailed protocol for these assays, please refer to Svoboda et al., 2016^35^.

### Cytokine cloning and expression

First, the amino acid sequence of zebrafish Kitlga/b, was subjected to protein structure prediction and hydrophobicity analysis using Phobius tool (http://phobius.sbc.su.se/). Two large hydrophobic regions (amino acid positions 1 to 24 and 206 to 224 for Kitlga, and amino acid positions 1 to 31 and 185 to 209 for Kitlgb) corresponding to the putative signal peptide (SP) and transmembrane (TM) domains, respectively, were identified (Supplemental Figure 1A). To generate a Kitlga/b version devoid of both domains, sequence specific primers (Supplemental Table 1) were used for PCR amplification of each Kitlg cDNA fragment corresponding to amino acid 25 to 182 (Kitlga) and 31 to 187 (Kitlgb) from adult zebrafish retina. We used the baculovirus expression system to produce soluble Kitlga/b in large quantities. We cloned the amplified fragment into modified pAc-GP67-B vector containing 6xHis and generated the recombinant baculovirus by co-transfection of pAc-His-Kitlga/b and BaculoGold Bright Baculovirus DNA into sf21 insect cells. Virus-infected cells expressed GFP (Supplemental Figure 1B) and secrete recombinant His-Kitlga/b extracellularly. Finally, we purified the secreted proteins on a Ni2+-NTA agarose column and used them for the following experiments (Supplemental Figure 1C). To generate Kitlga/b the construct for mRNA injection experiments, the full length cDNA fragment was amplified using RT-PCR using sequence specific primers (Supplemental Table 1). Erythropoietin protein and mRNA has been prepared as previously described^22,35,36^.

### Additional methods

For experimental details on the generation of recombinant cytokines, mRNA microinjection, benzidine staining, image analysis, qPCR, FACS-sorting, RNAseq and transcriptomics, please refer to supplemental methods.

### Data sharing statement

RNAseq data are available in ArrayExpress under accession number E-MTAB-8800. Plasmids for cytokine expression are available via Addgene with the accession numbers 140292 (pAc-His-zfKitlga) and 140293 (pAc-His-zfKitlgb).

## Results

### Kit ligands promote erythroid and myeloid expansion of whole kidney marrow cells

First, we cloned and expressed recombinant zebrafish Kitlga and Kitlgb. We amplified the mature form of zebrafish *kitlga* and *kitlgb* lacking the signal peptide-encoding region, intracellular and transmembrane region (Supplemental Figure 1A) from adult retina and we produced recombinant Kitlga and Kitlgb in sf21 insect cells (Supplemental Figure 1B, C). His-tagged purified proteins were used in the following experiments.

To test the biological activity of Kitlg proteins, we designed an experiment using whole kidney marrow cells isolated from adult, 6-month old zebrafish. To reveal differences that would suggest biological activity and potential cooperation of Kitlg proteins with other recombinant cytokines, we treated the cells with various factors, or their combination, and subsequently counted them at specific time points. After 3 days in culture, the potential enhancement in the myeloid lineage was assessed. We added both Kit ligands, either separately or combined with Gcsfa, which has been previously shown to support myeloid cell fate^37^. Interestingly, we observed an increase in the number of cells in all the conditions tested when compared to untreated control (Figure 1A). In agreement with previously published studies, Gcsfa promoted significantly the growth of whole kidney marrow cells, and the effect was further potentiated by the addition of either of the Kit ligands. This additional effect tended to be similar for both Kit ligand paralogs, however this cooperation proved to be statistically significant only for Kitlgb, but not Kitlga. The Kit ligands alone exhibited reproducible mild activity that was, however, statistically not significant. To verify that the addition of Gcsfa along with Kitlga or Kitlgb promoted myelopoiesis, we morphologically characterized cultured cells (Supplemental Figure 2A, top). At day 3, the culture mostly comprised of monocytes and macrophages. Although the addition of Kitlga or Kitlgb to Gcsfa increases cumulative number of cells (as shown in Figure 1A), the overall composition of cell culture and proportion of different cell types remained similar with or without the Kit ligands.

**Figure 1.**
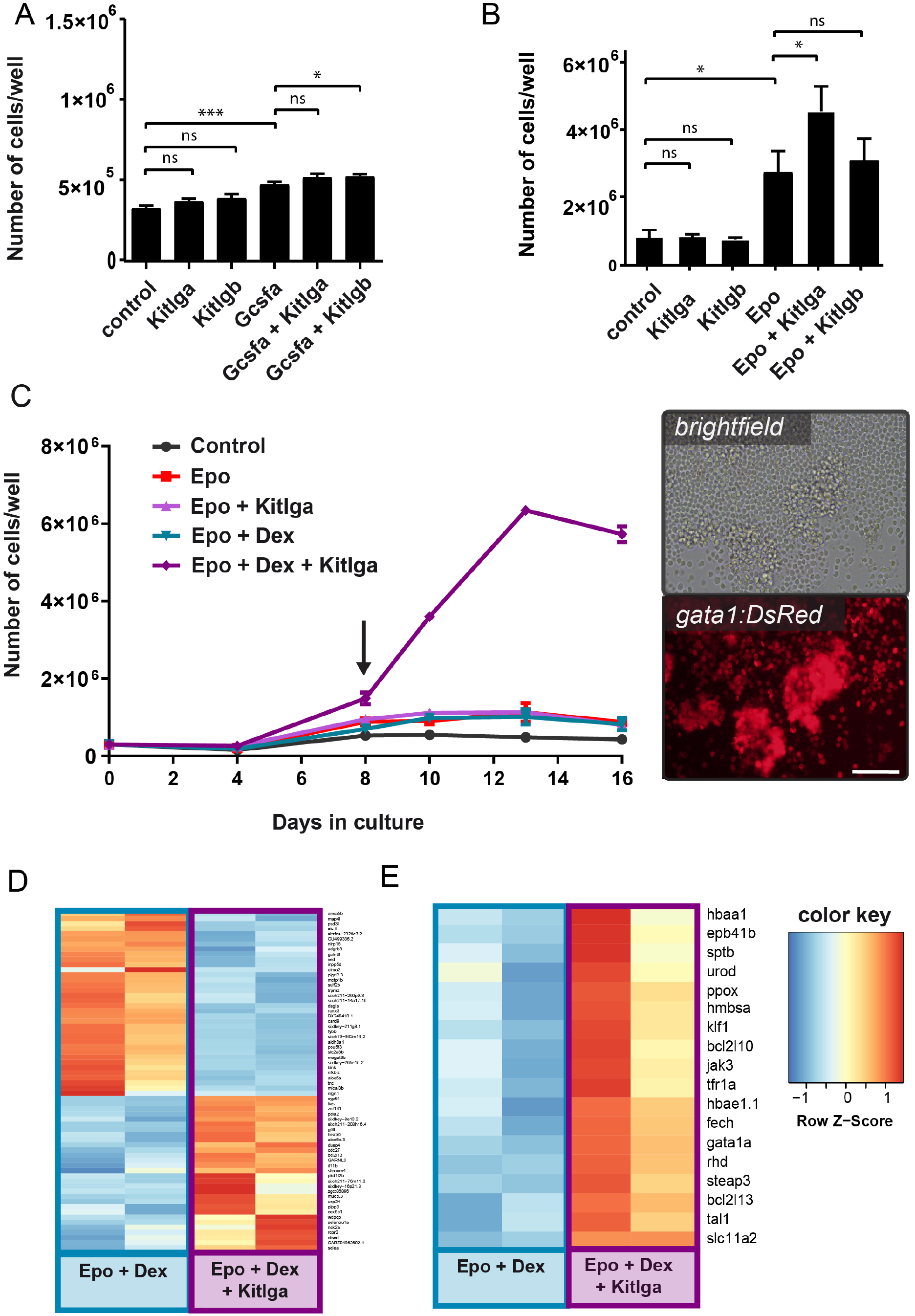
The effect of zebrafish Kitlga and Kitlgb on self-renewal and proliferation of hematopoietic cells *ex vivo*. (A, B) Quantification of the number of whole kidney marrow cells after treatment with PBS (control) or specific cytokine combinations for 3 days (A) and 7 days (B). Statistical significance was assessed using standard one-way ANOVA. n.s. = p>0.05, *p≤ 0.05, **p ≤ 0.01, ***p ≤ 0.001, ****p ≤ 0.0001. (C) Growth curve for *ex vivo* culture of whole kidney marrow cells with different cytokine combinations. Biological duplicates for every condition and timepoint are shown. Representative photomicrographs (right) of whole kidney marrow cells isolated from transgenic *gata1:DsRed* fish treated with Epo, Dex and Kitlga and cultivated for 8 days. Images were acquired on Olympus IX70 inverted microscope equipped with the Olympus DP72 camera using a 20x objective. Scale bar represents 100 um. (D, E) An expression heatmap of Epo, Dex and Epo, Dex, Kitlga treated cells showing the top predicted differentially expressed genes (D) and representative markers of erythroid differentiation (E). Color coding by z-score for heatmaps in E and F.

Next, we tested the activity of Kit ligands during erythroid cell expansion. Cells were treated with both Kitlga and Kitlgb, either separately or combined with Epo. The increase in cell number was quantified after 7 days in culture. In agreement with previously published studies^22,36^, Epo boosted cell growth significantly and again, there was a mild increase in the number of the cells when treated with any of the Kit ligands alone. Strikingly, when Epo was combined with Kitlga, we measured a significant increase in the number of cells at 7 days in culture compared to Epo alone (Figure 1B). Although, the additive effect of Kitlgb was not significant, the trend to increase the number of cells in the presence of Epo was still observed. To examine composition of cell culture, we further performed morphological characterization of cultured cells. At day 7, the culture mostly comprised of erythroid cells upon Epo stimulation as expected (Supplemental Figure 2A, bottom) and the addition of Kitlga led to an increase in number of erythroid progenitors, which explains elevated number of cells in this culture.

With the aim of maximum expansion of erythroid progenitors, we decided to improve the conditions for suspension culture of whole kidney marrow cells. Although there were no existing reports of suspension culture of zebrafish or hematopoietic progenitor cells, we supposed that similar conditions might be effective in suspension cultures as was shown in human and mouse ^10,38^. As different steroids were reported as important players in the maintenance of self-renewal and proliferation of erythroid progenitors in chicken, mouse and human ^38–41^, we included Dex and tested its different concentrations and combinations together with Epo and Kitlga, the more potent Kitlg paralogue.

We found that, in line with previous findings in other vertebrates, Kitlga, Epo and Dex act synergistically in *ex vivo* cultures, enabling erythroid expansion (Figure 1C, left). When compared to the cells treated with Epo and Dex, cells treated with Epo, Dex and Kitlga formed islets of round, highly *gata1:DsRed-*positive cells, corresponding to erythroid progenitors (Figure 1C, right). Additionally, we observed that Epo can cooperate with Kitlga to promote the growth of large erythroid colonies in colony forming assays in methylcellulose (Supplemental Figure 2B). When Kitlga was combined with Epo in kidney marrow culture from *gata1:DsRed* transgenic fish, we observed significantly more, highly *gata1*:DsRed positive colonies that were also significantly larger (Supplemental Figure 2C, D).

### Kitlga treatment, when combined with Epo and dexamethasone, leads to upregulation of erythroid-, translation- and cell cycle-related gene expression

To gain deeper insight into the mechanism of Kitlga action when combined with Epo and Dex in kidney marrow suspension cultures, we performed an RNAseq experiment, comparing samples from Epo, Dex, Kitlga and Epo, Dex-treated cells. Samples were collected after 8 days in culture. This was the first time-point where we could observe a significant increase in proliferation of erythroid cells in the presence of Epo, Dex and Kitlga, as shown in the growth curve (Figure 1C, left).

RNAseq analysis revealed that both culture conditions resulted in high expression of erythroid marker genes (data not shown). However, addition of Kitlga to the Epo, Dex combination caused an additional increase in expression of erythroid genes. Among the upregulated differentially expressed genes (see heatmap in Figure 1D and Supplemental Table 4), we identified several erythroid (*g6fl*, *lias*) and cell cycle (*cdc27*) specific genes, confirming additional Kitlga-mediated erythroid cell expansion over Epo and Dex treatment alone. On the contrary, downregulated genes were mostly linked to the myeloid cell lineage (*pigrl2*.3, *blnk*, *nfkbiz*, *alox5a*, *inpp5d*), providing further evidence that Epo, Dex, Kitlga-induced expansion led to a more enriched erythroid cell population as compared to Epo, Dex culture alone. Next, we examined the expression of both kit receptor and revealed that the expression of kita is predominant in adult erythroid progenitor cells (average raw counts 285 (kita) vs. 3 (kitb)).

Finally, we were interested specifically in changes in expression of erythroid-specific genes. The analysis of RNAseq data reveled that, compared to cells treated with Epo and Dex, cells treated with Epo, Dex and Kitlga display higher expression of many of the important erythroid genes – namely globins (*hbae1.1*, *hbaa1*), genes involved in heme and iron metabolism (e.g. *urod*, *slc11a2*, *steap3*), erythroid cytoskeleton (e.g. *rhd*, *sptb*, *eph41b*) and signaling (*gata1a*, *jak3*) (Figure 1E, Supplemental Table 5). Based on GO prediction, we revealed an enrichment in specific categories of biological processes – namely in translation (<u>GO:0006412</u>, <u>GO:0002181</u>, <u>GO:0000028</u>, <u>GO:0000027</u>), erythropoiesis (<u>GO:0042541</u>, <u>GO:0030218</u>, <u>GO:0048821</u>) and cell cycle regulation (<u>GO:0051726</u>). For a full list of enriched GO terms in all three standard GO term categories, refer to Supplemental Table 6.

### Both Kit ligands cooperate with Epo to promote erythroid expansion in zebrafish embryos

To test the function of both Kit ligands *in vivo*, we microinjected cytokine mRNA(s) into 1-cell stage embryos. When combined with *epo*, both *kitlga* and *kitlgb* showed an enhancement in *lcr:EGFP* expression (Figure 2A) and hemoglobinization (Figure 2D) at 72 hpf compared to *epo* alone. Quantification of *lcr:EGFP* positive cells in the tail region (Figure 2B), FACS analysis of total *lcr:EGFP* positive cells (Figure 2C and Supplemental Figure 3) as well as benzidine staining in the tails (Figure 2E) of 72 hpf embryos confirmed the observed trend. Note that even though similar trend to increase number of erythroid cells, in combinations with *epo*, was observed for both Kit ligands under all three experimental readouts, the effect was more robust and statistically significant only for *kitlgb* in all types of experiments. Next, quantitative PCR (qPCR) analysis from whole injected embryos also showed an increase in the expression of erythroid markers at 72 hpf (Figure 2F) when *epo* was combined with either of the ligands, however, these changes were rather subtle. Finally, in agreement with previous studies, we observed increased number of melanocytes after the overexpression of Kitlga but not Kitlgb.

**Figure 2.**
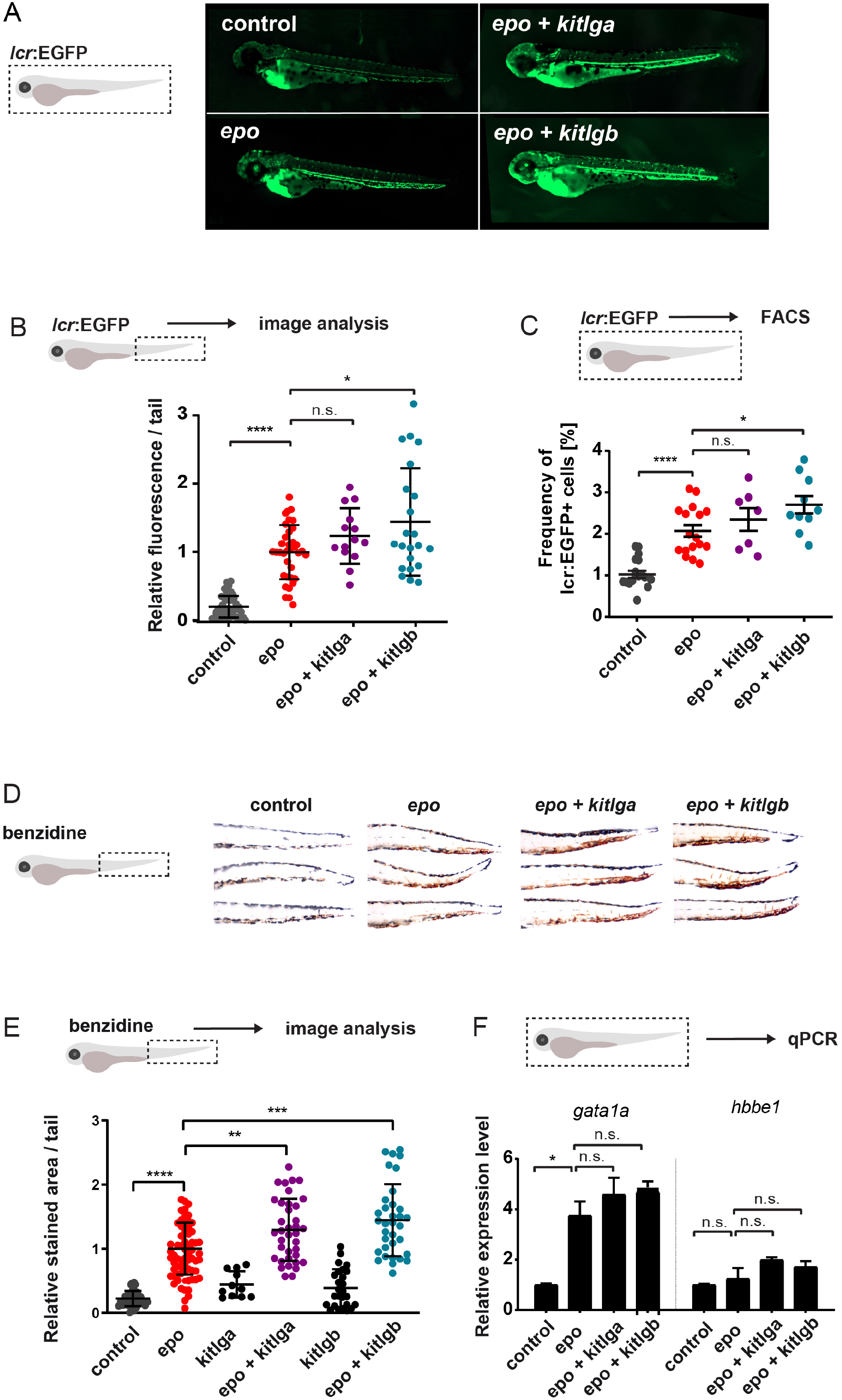
*In vivo* effects of *kitlga* and *kitlgb* at 72 hpf. (A) Representative micrographs of *lcr:EGFP* transgenic reporter larvae at 72 hpf after injection of corresponding mRNAs. Control represents uninjected larvae. Images were acquired on Zeiss Axio Zoom.V16 with Zeiss Axiocam 506 mono camera and ZEN Blue software. (B) Quantification of the *lcr:EGFP* positive cells in embryonic tails from fish in (A). Values were plotted using the mean of *epo* as the relative expression fold 1. (C) FACS analysis of *lcr:EGFP* positive cells in whole embryos. (D) Representative micrographs of benzidine stained embryonic tails at 72 hpf after injection of corresponding mRNAs. Control represents uninjected embryos. NOTE increase in pigmentation in the dorsal part of *kitlga*-injected embryos. Z-stacks were acquired on Zeiss Axio Zoom.V16 with Zeiss Axiocam 105 color camera and processed using Extended Depth of Focus module in the ZEN blue software. (E) Quantification of the benzidine stained embryonic tails in (D). Values were plotted using the mean of *epo* as the relative expression fold 1. (F) qPCR expression analysis of *gata1a* and *hbbe1* from whole injected embryos at 72 hpf. Data have been normalized using *mob4* as the housekeeping gene and using uninjected control as the relative expression fold 1. Bars represent a mean of three triplicates with SD. n.s. = p>0.05, *p≤ 0.05, **p ≤ 0.01, ***p ≤ 0.001, ****p ≤ 0.0001.

### The expression of both kit receptors is gradually increasing during embryonic development

To better understand the mechanism of cooperation of Epo with Kit ligands, we first analyzed the expression of both kit receptors in tissues and during the development by qPCR. Among all the tissues analyzed, the highest *kita* expression was observed in kidney (Supplemental Figure 4A) while the highest kitb expression was observed in the retina (Supplemental Figure 4B). During zebrafish embryonic development, the highest relative expression was observed at 7 dpf (*kita*) and 72 hpf (*kitb*) (Figure 3A). As we saw a significant expansion of erythroid, *gata1:DsRed*+ cells in *ex vivo* cultures, we aimed to analyze the dynamics of the receptor expression in erythroid cells during development. Therefore, we collected samples from 24 hpf, 48 hpf and 72 hpf *gata1:DsRed* and *lcr:EGFP-*embryos and FACS-sorted *gata1*:*DsRed*-positive cells (Figure 3B) or *lcr:EGFP-*positive cells (Figure 3C), respectively. In both cases, we found that there is a gradually increasing expression of both receptors in erythroid cells during the course of early development.

**Figure 3.**
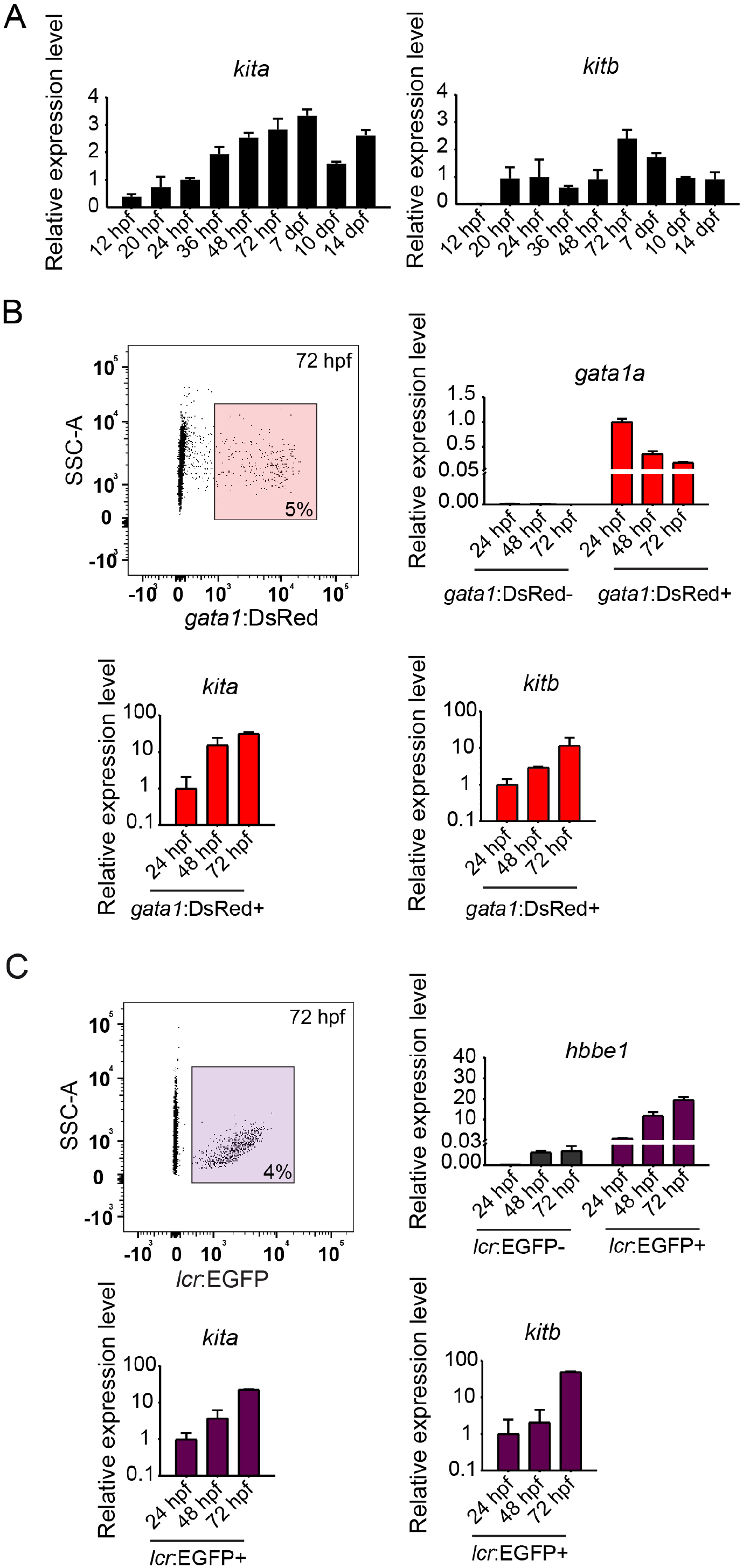
Expression of *kit* receptors in the development. (A) qPCR expression analysis of *kita* (left) and *kitb* (right) at different developmental stages. Data was normalized using *ef1a* as the housekeeping gene and using the 24 hpf sample as relative expression fold 1. (B) Sorting strategy used for FACS isolation of *gata1:DsRed* positive cells (top left) from transgenic zebrafish at 24 hpf stage. qPCR expression analysis of *gata1* (top right), *kita* (bottom left) and *kitb* (bottom right) in *gata1:DsRed* negative and positive cells. Data was normalized using *ef1a* as the housekeeping gene and using the *gata1:DsRed* positive 24 hpf sample as relative expression fold 1. (C) Sorting strategy used for FACS isolation of *lcr:EGFP* positive cells (top left) from transgenic zebrafish at 24 hpf stage. qPCR expression analysis of *gata1* (top right), *kita* (bottom left) and *kitb* (bottom right) in *lcr:EGFP* negative and positive cells. Data was normalized using *ef1a* as the housekeeping gene and using the *lcr:EGFP* positive 24 hpf sample as relative expression fold 1.

### Kita mediates Kitlg-dependent erythroid expansion in vivo

To understand the molecular basis of the effect we observed on erythroid cells, we used the *sparse* mutant (*kita*^*b*5/b5^) zebrafish, lacking the functional Kita receptor. We injected mRNA for *epo*, *kitlga* and *kitlgb*. We observed that in *sparse* mutants, the number of erythrocytes is slightly decreased in *epo* injected embryos and the effect of cooperation between any of the *kitlg* and *epo* is completely lost (Figure 4A). These findings are also supported by qPCR data showing that the expression of β-globin (*hbbe1*) and *gata1a* is not increased after the injection of *epo* with either of the kit ligands, as is the case in wild-type embryos (Figure 4B).

**Figure 4.**
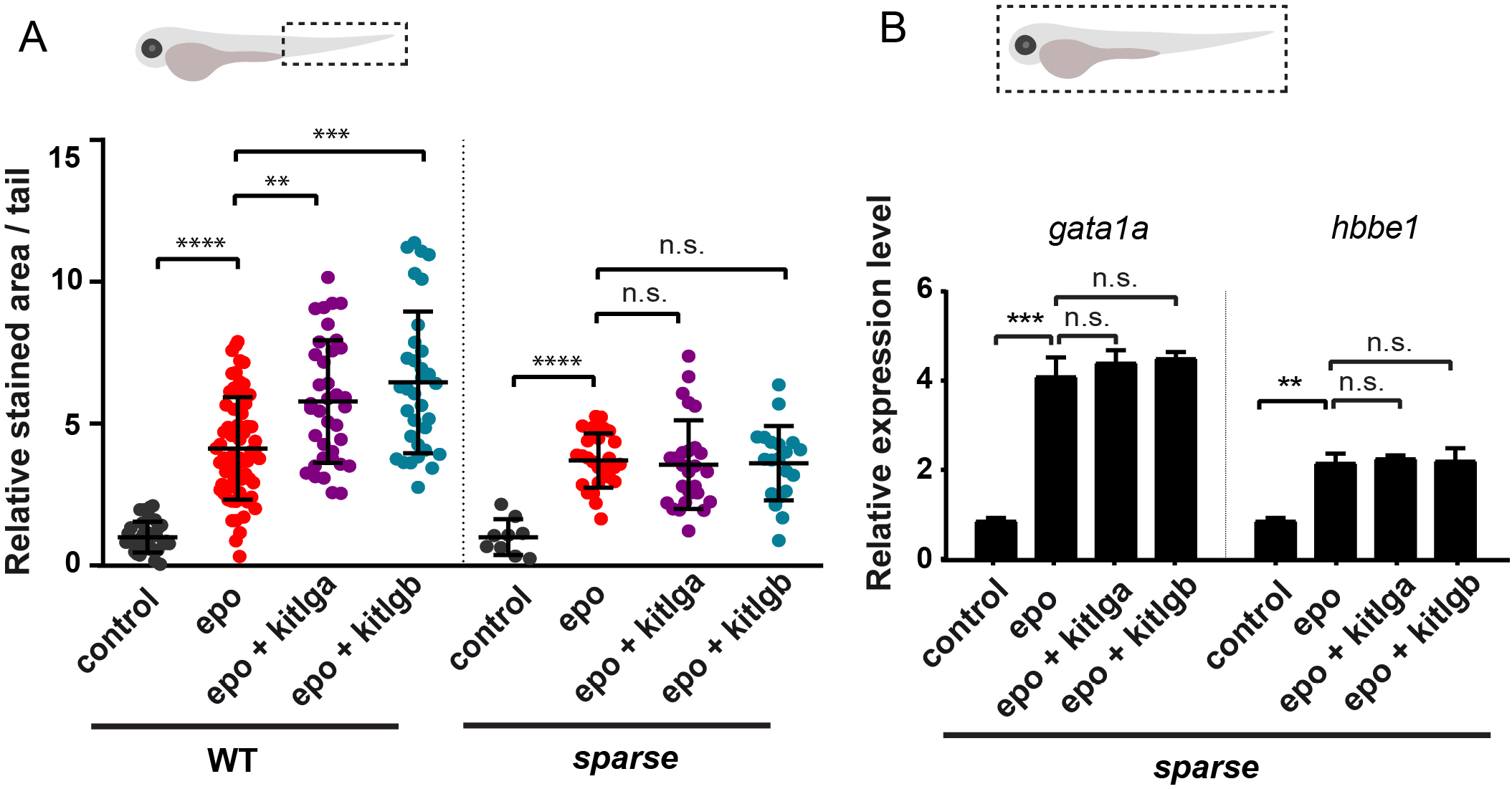
The effect of *kitlga* and *kitlgb* in *sparse* mutants. (A) Quantification of the benzidine staining in the tail region of control (uninjected) or *epo*, *kitlga* and *kitlgb* mRNA injected wild-type or *kita* mutant embryos (*sparse*). Values were plotted relatively to the mean of *epo* (=1). NOTE that for demonstrative purposes, Figure 4A left is re-used from Figure 2C. (B) qPCR expression analysis of *gata1a* and *hbbe1* from whole injected *kita* mutant embryos (*sparse*) at 72 hpf. Data was normalized using *mob4* as the housekeeping gene and uninjected control as the relative expression fold 1. Bars represent mean of three triplicates with SD. n.s. = p>0.05, *p≤ 0.05, **p ≤ 0.01, ***p ≤ 0.001, ****p ≤ 0.0001.

To summarize, we have cloned and expressed both Kit ligand zebrafish paralogs. Using the purified recombinant proteins, we tested various cytokine combinations in suspension culture of whole kidney marrow cells and defined an optimal factor combination, composed of Epo, Kitlga and dexamethasone, which enables efficient expansion of erythroid progenitors. Also, we tested gain of function effect of both *kitlga and kitlgb* in embryonic hematopoiesis and observed an increase in the number of erythroid cells in *epo, kitlgb* and *epo, kitlgb* mRNA-injected fish compared to *epo* alone. Finally, by employing the *kita* (*sparse*) mutants, we showed that Kita is mediating Kitlg-dependent erythroid expansion *in vivo*.

## Discussion

In this study, we explored the role of zebrafish Kit ligands in hematopoiesis. In zebrafish, multiple paralogs of many important genes resulted from an extra round of whole genome duplication 250-350 million years ago^42,43^, including hematopoietic cytokines. After such an event, the process of pseudogenization often leads to a loss of function of one of the paralogs^44–46^. Alternatively, paralogs of an ancestral gene can either remain functionally redundant, split the original function between the two (subfunctionalization), or acquire completely new functions (neofunctionalization)^47^. Here, we studied the hematopoietic function of two retained zebrafish paralogs of a crucial hematopoietic cytokine, the Kit ligand (KITL or Stem Cell Factor – SCF), designated as Kitlga and Kitlgb.

So far, KIT signaling has not been associated with erythropoiesis in zebrafish^24,25^. This is in contrast with previous studies in human^8^, mouse^4^ and chicken^9,16,48^, where it has been shown that KITL is critical for proper erythroid development^5,15,49^. By focusing on Kit function during erythropoiesis in zebrafish, we were able to prove for the first time, using *ex vivo* and *in vivo* gain- and loss-of-function experiments, that Kit signaling does indeed play a role in zebrafish erythroid differentiation. Based on RNAseq and GO analysis, this likely happens in a similar manner as in other vertebrate species via induction of cell proliferation coupled with differentiation^40,48^, promoting cell cycle progression and enhancing protein translation in differentiating cells.

To examine the role of Kit ligands *ex vivo*, we used clonal assays and liquid culture experiments^23,36,37,50^. Since the reported cross-reactivity of mammalian hematopoietic cytokines in zebrafish is very limited^35^, it was essential to use *bona fide* zebrafish Kit ligands as recombinant proteins. Similarly as described for other cytokines^7,9,35,36^, we cloned the extracellular portion of the corresponding cDNAs, lacking the N-terminal signal peptide, transmembrane domain and cytosolic part of both cytokines. This approach allowed us to generate soluble form of Kit ligands to stimulate kidney marrow progenitor cells *ex vivo*^8,13^.

As a result, treatment of hematopoietic progenitor cells isolated from kidney marrow with Kitlga, in combination with the other master regulator of erythroid lineage commitment, Epo^22,23^, enabled erythroid expansion of cells in suspension cultures and clonal assays, as in other vertebrates^1,8,10,11^. Such an effect was observed also for Kitlgb, however its cooperation with Epo was less pronounced and was not statistically significant. We hypothesize that soluble form of Kitlgb that lacks the transmembrane and cytosolic domain is less effective to stimulate differentiation of hematopoietic progenitors towards erythroid lineage and that the main role of Kitlgb is stimulation of neighboring cells in stem cell niches via contact dependent signaling.

Next, we also tested the activity of Dex in addition to Epo and Kitlga, the more effective soluble Kitlg paralogue. Dex is a glucocorticoid receptor agonist that has been reported to support self-renewal and differentiation of erythroid progenitors^10,38–40^ and indeed, addition of Dex to Kitlga and Epo led to synergistic erythroid expansion in suspension cultures.

To understand the mechanism triggering this substantial cell expansion, we performed an RNAseq experiment. We decided to compare the effect of Epo and Dex versus Epo, Dex, and Kitlga treatment on adult kidney marrow cells. Transcriptional profiling showed expression of erythroid specific genes in both conditions, as expected due to Epo-mediated signaling. However, we observed relatively subtle changes in expression between the Epo and Dex and the Epo, Dex, and Kitlga cultures. The trends of expression changes showed a distinct erythroid fingerprint in cells treated with Epo, Dex and Kitlga, compared to a more mixed/myeloid one in cells treated with Epo and Dex alone, as suggested by the increased expression of myeloid-specific genes. In addition, we found that the prominent receptor expressed in adult erythroid progenitor cells is kita, indicating that the effect of erythroid expansion is mediated by this receptor. This corresponds to the fact that isolation of kidney marrow cells yields, in addition to progenitor cells, also a fraction of myeloid cells with the ability to survive in the culture. However, this effect becomes negligible during erythroid expansion of cells treated with Epo, Dex, and Kitlga, when myeloid cells are overgrown by expanding erythroid progenitors. Importantly, gene ontology analysis revealed an enrichment especially in translation and expression of erythroid-specific genes in Epo, Dex and Kitlga cultures. Therefore, we hypothesize that Kitlga might enhance erythroid cell development and cell proliferation potentially through upregulation of genes related to protein translation and cell cycle progression.

To confirm these findings, we decided to test, whether Kit signaling is involved also in zebrafish erythropoiesis *in vivo* using full length form of both kit ligands. For *ex vivo* experiments, we used soluble forms, whereas for *in vivo* experiments, we used the full-length coding sequence, including transmembrane and cytosolic domains, as used in previous reports^27^. It has been shown that the function of soluble and transmembrane forms under physiological conditions is different^13^. The soluble form of the protein is distributed throughout the whole organism via circulation and affect distant tissues. It has been shown that this form is required and sufficient for proper erythroid signaling in cultured cells^16^. On the other hand, according to previous studies, the major site of action of the transmembrane form of Kit ligand might be in stem cell niches^13^. We hypothesize that full length Kitlgb can contribute, in addition to HSPCs^27^, also to erythroid expansion *in vivo*. As a result, we found that the full length *kitlga* was potent in expansion of erythroid cells at 72 hpf when injected together with *epo* mRNA. Interestingly, similarly to *kitlga*, also the *kitlgb* was able to mediate such erythroid expansion, when injected *in vivo* in full-length form, but this effect was markedly decreased in *ex vivo* experiments using short soluble form. This supports our hypothesis that there is a difference in potency between soluble and full-length form of Kitlgb. To prove this, we generated shortened *kitlga* and *kitlgb* mRNA, corresponding to the soluble region of Kitlg proteins and indeed, there was a decrease in a potency of soluble *kitlgb* to stimulate erythropoiesis (data not shown), whereas soluble *kitlga* remained equally active.

Finally, we investigated, if Kita receptor is dispensable in Kitlg-mediated erythroid cell expansion *in vivo*. As a result, we did not observe the cooperation of *epo* with either of the *kit* ligands in erythroid differentiation in *kita* (*sparse*) mutant fish. These data suggest that Kita is responsible for Kit signal transduction in erythroid cells. Although Kitb might also be involved in this process, it couldn’t compensate for missing Kita, and Kita seems to be required for Kit signaling in zebrafish erythroid cells. In addition to this, we also observed decreased number of erythroid cells in both, control and in *epo* injected, *sparse* mutants when compared to the wild-type embryos. This indicates that endogenous Kit signaling plays role also under physiological conditions during the normal erythropoiesis.

To summarize, in this study we have addressed the question of the role of Kitlg in hematopoiesis in zebrafish using both *in vivo* and *ex vivo* approaches. Importantly, we established optimized liquid culture conditions for erythroid expansion of kidney marrow progenitor cells that require the addition of Kitlga. Together with colony forming assays, suspension cultures might serve as indispensable tools in disease modelling, allowing analysis of normal and aberrant hematopoiesis in zebrafish blood mutants (*vlade*, *moonshine*, *mindbomb* etc.)^51^. This will enable further biochemical, genomic and proteomic analysis of *ex vivo* cultured and expanded cells.

Interestingly, zebrafish Kitlga and Kitlgb seem to possess only partial redundancy in its function: Kitlga retains the role in melanogenesis and pigmentation, which is lost in the case of Kitlgb, but both ligands potentially contribute to the overall process of erythroid commitment and development in zebrafish *in vivo*. Although this function is not required in steady state erythroid development, it might become important in stress conditions. The exact mechanism of functional diversification of Kit ligands has yet to be investigated but altogether, we demonstrate that role of Kit signaling is evolutionarily conserved with respect to erythroid development as previously unnoticed^24^, and therefore it strengthens the use of zebrafish as a model to study normal and aberrant human hematopoiesis.

## Acknowledgements

We thank Nikol Pavlu,Tereza Hojerova and Tereza Hingarova for animal care, Tereza Mikulasova and Martina Hason, for graphical work and cDNA samples, Trevor Epp for editing the manuscript and Leonard Zon for providing *gata1*:DsRed and *lcr*:EGFP reporter fish lines. Supported by the Czech Science Foundation (16-21024S), Ministry of Health (NV19-07-00412) and 68378050-KAV-NPUI to PB. OS was partially funded by American Heart Association (19POST34380328).

## Authorship contribution

J.O., O.M., O.S. performed the experiments, J.O, O.S., P.S., D.T., M.K., and P.B. designed the experiments, analyzed the data and wrote the manuscript.

## Conflict of interest

The authors declare no conflict of interest.

## Notes

### Competing Interest Statement

The authors have declared no competing interest.

