## Supplementary information for "Zebrafish Kit ligands cooperate with erythropoietin to promote erythroid cell expansion"

### Supplemental methods

#### mRNA microinjections

mRNAs were transcribed *in vitro* using mMessage mMachine SP6 kit (Roche) from a linearized pCS2+ vector containing PCR-amplified product and purified by lithium chloride extraction. 1-cell stage embryos were injected with 2 nl of the injection mix. The concentration of mRNA was 300 ng/ul for each cytokine, which equals to 600 pg of each mRNA per embryo.

#### Benzidine staining and image analysis

72 hpf embryos were anesthetized by Tricaine (MS-222) and incubated for 5 minutes on ice. Next, embryos were washed with cold PBS and fresh benzidine solution composed of o-dianisidine (Sigma-Aldrich, D9143) (0.6 mg/ml), sodium acetate pH 4.5 (0.01 M), hydrogen peroxide (0.65%) and ethanol (40%) was added. Fish were incubated for 10 mins in dark, followed by wash with PBS. Fish were imaged using Zeiss Axio Zoom.V16 with Plan Neofluar Z 1.0x objective (total magnification 125x) and Zeiss Axiocam 105 color camera. Acquired brightfield z-stacks were processed using the Extended Depth of Focus module in ZEN blue 2.3 software and images were quantified in Fiji using the Color Threshold and Analyze particles features of the software.

#### Fluorescence imaging and quantification

72 hpf embryos were anesthetized using Tricaine (MS-222) and incubated for 5 minutes on ice. Fish were imaged using Zeiss Axio Zoom.V16 with Plan Neofluar Z 1.0x objective (total magnification 125x) and Zeiss Axiocam 506 mono camera. Acquired brightfield z-stacks were quantified in Fiji using the Threshold and Analyze particles features of the software.

#### Preparation of embryonic hematopoietic cells for FACS

24 hpf, 48 hpf and 72 hpf embryos were treated with Liberase TM (Roche, 05401119001), diluted 100x in PBS and incubated at 37°C for 1, 1.5 or 2.5 hours, respectively for each stage. Suspension was filtered using a 30µm filter, centrifuged and resuspended in PBS.

**RNA isolation from embryos and qPCR**

RNA from whole injected embryos was isolated using PureLink Micro Kit (Invitrogen, 12183-016) according to the manufacturer's instruction that include DNase I treatment. Alternatively, sorted cells were collected into TRI Reagent (Sigma, T3934) and RNA was isolated with RNeasy Plus Mini Kit (Qiagen, 74136). cDNA was synthesized using iScript™ cDNA Synthesis Kit (Bio-Rad). qPCR analysis was performed in triplicate with the LightCycler 480 and SYBR Green I (Roche). *ef1a* or *mob4* housekeeping genes were used to normalize the RNA content of samples, and data were calculated with the ΔΔCt method. The qPCR primers are listed in Supplemental Table 2.

**Cytology**

Hematopoietic cells were removed from cultures and concentrated by centrifugation at 300g for 2 minutes before being smeared onto glass slides. Air-dried smears were fixed and stained with May-Grünwald Giemsa (Sigma) and images were acquired with a Leica DM 2000 microscope and Zeiss Axiocam 105 color camera.

**RNAseq and transcriptomics**

Total RNA was isolated from 6-7×$10^6$ WKM cells cultivated in the presence of Epo, Dex or Epo, Dex and Kitlga in biological duplicates, according to the PureLink Micro Kit (Invitrogen) manufacturer's protocol that includes DNase I treatment. The quantity of isolated RNA was measured spectrophotometrically using NanoDrop ND-1000 (Thermo Fisher Scientific) and its quality was analyzed by Agilent 2100 Bioanalyser (Agilent Technologies). RNA integrity number, which is regarded as criteria for high-quality total RNA, ranged between 9.5 and 10. Takara SMARTer Stranded Total RNA-Seq Kit v2 (Takara Bio) was used for cDNA library preparation starting with 3.5 ng of total RNA. Library size distribution was evaluated on the Agilent 2100 Bioanalyzer using the High Sensitivity DNA Kit (Agilent Technologies). Libraries were sequenced on the Illumina NextSeq® 500 instrument (Illumina) using 75bp single-end configuration yielding on average 47 million reads per sample.

Read quality was assessed by FastQC[1.] Individual steps of read processing were removing of sequencing adaptors with Trimmomatic (v0.36)[1], removing of ribosomal RNA with SortMeRna (v2.1)[2], mapping to reference transcriptome GRCz11[3] (Ensembl assembly version 98) and quantifying with Salmon[4].

Aggregation of transcript-level quantification to the gene level was performed with tximport[5] and served as input for differential expression analysis using DESeq2 R (v3.6.0) Bioconductor (v3.9) package[6]. Prior to the analysis, genes not expressed in at least two samples were discarded. Shrunken log2-fold changes using the adaptive shrinkage estimator[7] were used for differential expression analysis. Genes exhibiting minimal absolute log2-fold change value of 1 (for the genes that were upregulated in Epo, Dex, Kitlga condition) or -2.32 (for the genes that were downregulated in Epo, Dex, Kitlga condition) and statistical significance (adjusted p-value) of <0.1 between compared groups of samples were considered as differentially expressed for subsequent interpretation and visualization. Gene ontology (GO) enrichment analysis was done using gene length bias aware algorithm implemented in goseq R Bioconductor package[8]. The thresholds for up and down regulated genes entering the GO term analysis were log2-fold change > 0.415 and adjusted p-value < 0.1. Supplementary Table S6 shows GO terms with the cutoff of log2-fold change > 0.3 and p<0.0001 sorted by Odds ratio.

**References**


1 *http://www.bioinformatics.babraham.ac.uk/projects/fastqc*.

2 *https://bioinfo.lifl.fr/RNA/sortmerna/*.

3 *http://www.ensembl.org/Danio_rerio/Info/Annotation*.

4 Patro, R., Duggal, G., Love, M. I., Irizarry, R. A. & Kingsford, C. Salmon provides fast and bias-aware quantification of transcript expression. *Nature Methods* **14**, 417-419, doi:10.1038/nmeth.4197 (2017).

5 Soneson, C., Love, M. I. & Robinson, M. D. Differential analyses for RNA-seq: transcript-level estimates improve gene-level inferences. *F1000Res* **4**, 1521, doi:10.12688/f1000research.7563.2 (2015).

6 Love, M. I., Huber, W. & Anders, S. Moderated estimation of fold change and dispersion for RNA-seq data with DESeq2. *Genome Biol* **15**, 550, doi:10.1186/s13059-014-0550-8 (2014).

7 Stephens, M. False discovery rates: a new deal. *Biostatistics* **18**, 275-294, doi:10.1093/biostatistics/kxw041 (2017).

8 Young, M. D., Wakefield, M. J., Smyth, G. K. & Oshlack, A. Gene ontology analysis for RNA-seq: accounting for selection bias. *Genome Biol* **11**, R14, doi:10.1186/gb-2010-11-2-r14 (2010).

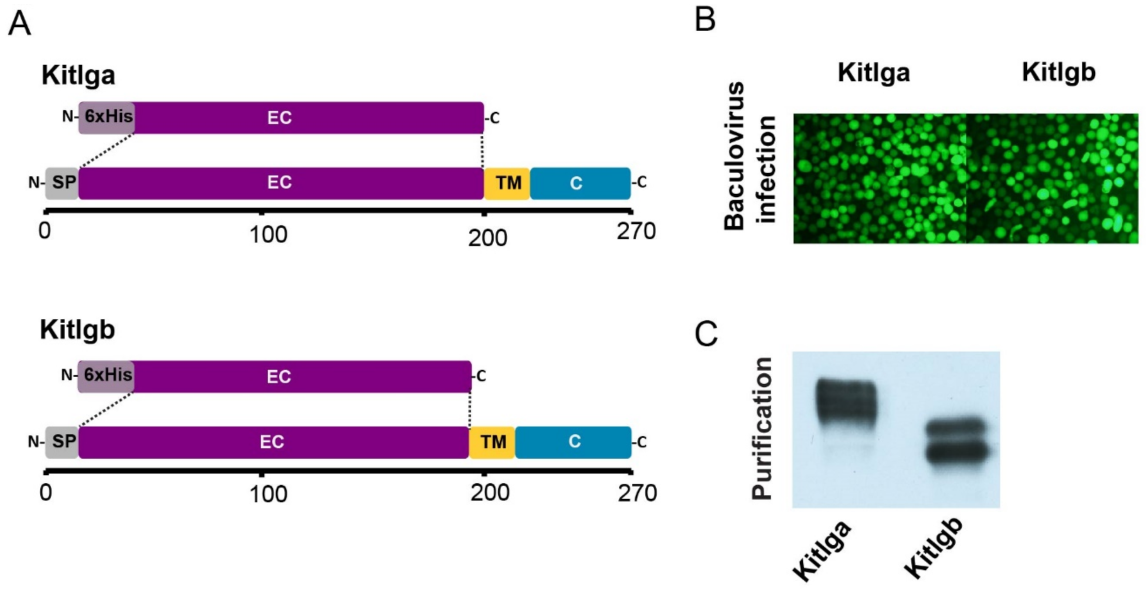


**Figure S1. Cloning and expression of zebrafish Kitlga and Kitlgb.**

(A) Sequence encoding the mature extracellular form (without the signaling peptide, transmembrane domain and cytosolic part) of each of the Kit ligands was amplified using RT-PCR from adult zebrafish retina and cloned into modified pAc-GP67-B vector containing 6xHis to generate recombinant His-tagged Kitlga and kitlgb proteins. SP - signaling peptide, EC – extracellular part, TM – transmembrane domain, C – cytosolic part, 6xHis - 6x histidine tag. Scale represents the number of amino acids.

(B) sf21 insect cells were co-transfected with BaculoGold Bright DNA (BD Biosciences) and Kitlga/b transfer vector leading to the production of a recombinant virus. Virus-infected cells were GFP-positive (top).

(C) Recombinant proteins were secreted from infected cells, purified using affinity chromatography and detected using anti-His antibody (bottom). Note, that recombinant proteins form multiple bands due to different states of glycosylation in insect cells.

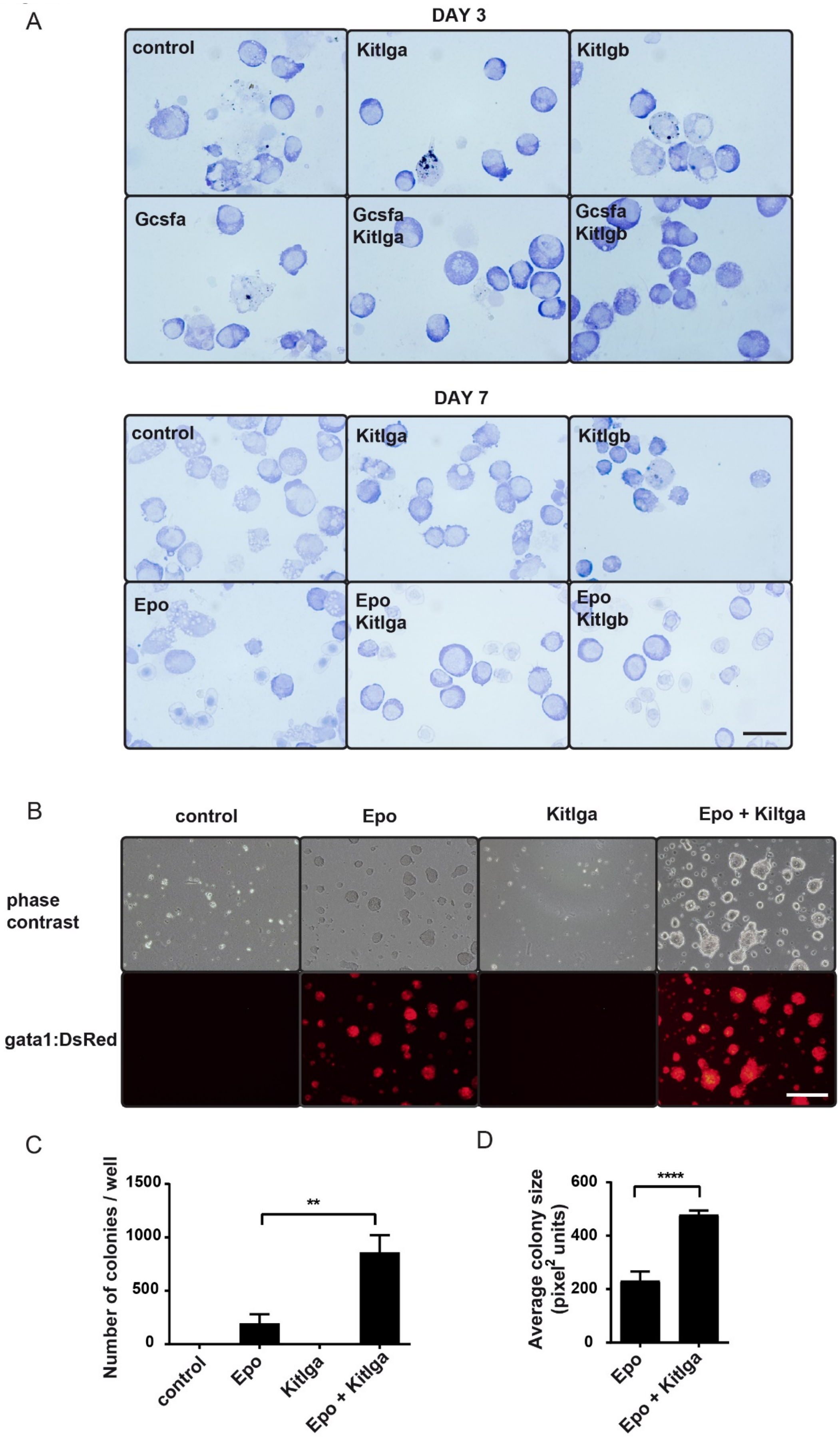
A
DAY 3
control
Kitlga
Kitlgb
Gcsfa
Gcsfa
Kitlga
Gcsfa
Kitlgb
DAY 7
control
Kitlga
Kitlgb
Epo
Epo
Kitlga
Epo
Kitlgb
B
control
Epo
Kitlga
Epo + Kiltga
phase
contrast
gata1:DsRed
C
Number of colonies / well
1500
1000
500
0
**
control
Epo
Kitlga
Epo + Kitlga
D
Average colony size
(pixel2 units)
600
400
200
0
****
Epo
Epo + Kitlga

**Figure S2. The effect of Kitlga in colony forming assays.**

(A) Representative fields of smeared cells cultured for three or seven days in the presence of specific cytokines or their combination. Cells were stained with May-Grünwald Giemsa. Top panel shows myeloid cells co-treated with Gcsf for three days and bottom panel shows erythroid cells that were co-treated with Epo for seven days in addition to Kit ligands. Scale represents 10 µm.

(B) Representative brightfield (top) and fluorescent (bottom) micrograph of *gata1*:DsRed positive colonies for each cytokine and their combination. Images were acquired on Olympus IX70 inverted microscope equipped with the Olympus DP72 camera, using a 10x objective. Scale bar represents 200 um.

(C) Quantification of the number of colonies that formed from whole kidney marrow cells after treatment with PBS (control) or specific cytokine combinations for 7 days. Each condition was counted for three separate wells of a 12-well plate from images acquired on Zeiss Axio Zoom.V16. Each bar represents mean with SD. Statistical significance was determined by one-way ANOVA.

(D) Quantification of the difference in the size of the colonies in (A). Each bar represents a triplicate mean with SD. Statistical significance has been determined by t-test.

n.s. = $p>0.05$, ***** $p \leq 0.05$, ****** $p \leq 0.01$, ******* $p \leq 0.001$, ******** $p \leq 0.0001$.

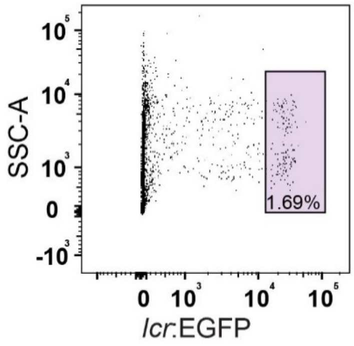


**Figure S3. Gating strategy used for quantification of *lcr*:EGFP cells**

Gating strategy employed during FACS analysis of *lcr*:EGFP cells isolated from 72hpf embryos in Figure 2C.

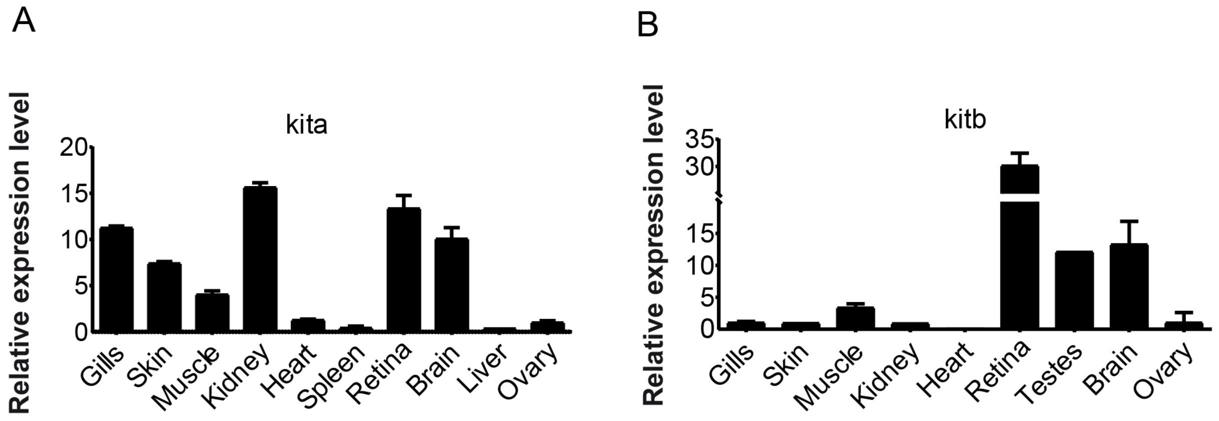


**Figure S4. Expression of kit receptors in tissues.**

qPCR expression analysis of *kita* (A) and *kitb* (B) in various adult tissues. Data were normalized using ef1a as the housekeeping gene and using ovary sample as relative expression fold 1.

**Supplemental Table 1. Primers for cloning kit ligands.**

| | Forward primer | Reverse primer |
|---|---|---|
| ***Protein*** | | |
| ***kitlga*** | CGGGATCCATTGAAATAGGAAATCCCAT | GCGGATCCTTACTCATTTGTACTATGTTGCGC |
| ***kitlgb*** | GCGGATCCGGGAGCCCTTTAACAGATGA | GCGGATCCTTAATGAACCGCAGAGTTCATGCC |
| ***mRNA*** | | |
| ***kitlga*** | GCGGATCCGGTTTCGCTGACATTGGAGT | GCGGATCCTTTGTTCCTGTAGGTTGGGC |
| ***kitlgb*** | GCGGATCCGGAGACACGGCTGATTTGTT | GCGGATCCATCCCGTTCTGGATATTCCC |

**Supplemental Table 2. Primers for qPCR.**

| | Forward primer | Reverse primer |
|---|---|---|
| ***gata1a*** | GTTTACGGCCCTTCTCCACA | CACATTCACGAGCCTCAGGT |
| ***hbbe1*** | CTTGACCATCGTTGTTG | GATGAATTTCTGGAAAGC |
| ***kita*** | CTATGTTGTCAAAGGCAATGCT | CCAGACGTCACTCTCAAAGGT |
| ***kitb*** | GGATACAGAATGAGTGAGCCTGA | CTCCAGCACCATCTCATCAC |
| ***efa1*** | GAGAAGTTCGAGAAGGAAGC | CGTAGTATTTGCTGGTCTCG |
| ***mob4*** | CACCCGTTTCGTGATGAAGTACAA | GTTAAGCAGGATTTACAATGGAG |

**Supplemental Table 3. Composition of zebrafish S13 medium.**

| zf S13 medium | final concentration |
|---|---|
| DMEM (Gibco, cat. no. 41966-029) | |
| Distilled Water (Gibco, cat. no. 15230-089) | |
| FBS, embryonic-stem-cell qualified (Biosera, cat. no. FB-1001S/500) | 10% |
| Carp Serum | 2% |
| BSA 10% (StemCell Technologies, cat. no. 09300) | 0.5% |
| NaHCO3, 7.5% | 0.2% |
| beta-Mercaptoethanol (Sigma-Aldrich, cat. no. M6250) | 0.1 mM |
| L-Glutamine, 0.2 M (Gibco, cat. no. 25030-081) | 4 mM |
| Hypoxanthine (Sigma-Aldrich, cat. no. H9636) | 200mg/100ml |
| FE-SIH iron supplement, 1000x (Sigma-Aldrich, cat. no. I3153) | |
| Penicillin-streptomycin, 100× (Gibco, cat. no. 15140122) | |
| solution A (1000x) | |
| solution B (50x) | |

| solution B in water | final concentration |
|---|---|
| L-Methionine | 10 mM |
| L-Phenylalanine | 20 mM |
| L-Alanine | 10 mM |
| Glycine | 50 mM |
| L-Threonine | 40 mM |
| L-Isoleucine | 40 mM |
| L-Proline | 10 mM |
| L-Valine | 40 mM |
| L-Aspartic Acid | 10 mM |
| L-Glutamic Acid | 25 mM |

| solution A in water | final concentration |
|---|---|
| Biotin (Sigma B4639) | 0.1% |

**Supplemental Table 4. Statistics of differentially expressed genes in cells treated with Epo, Dex, Kitlga compared to cells treated with Epo, Dex only.**

| Ensembl Accession ID | symbol | $log_2$FoldChange | p-adjusted |
|---|---|---|---|
| ENSDARG00000114387 | CU499336.2 | -2.46 | 1.40E-25 |
| ENSDARG00000098680 | si:zfos-2326c3.2 | -3.61 | 6.26E-23 |
| ENSDARG00000044694 | fybb | -2.36 | 4.25E-19 |
| ENSDARG00000069542 | si:dkey-8e10.2 | 3.01 | 5.19E-17 |
| ENSDARG00000042722 | blnk | -4.77 | 9.04E-16 |
| ENSDARG00000101641 | trpm2 | -2.35 | 1.77E-14 |
| ENSDARG00000102389 | ved | -2.50 | 6.59E-14 |
| ENSDARG00000060871 | mctp1b | -3.31 | 6.27E-13 |
| ENSDARG00000058476 | stc1l | -3.10 | 3.74E-12 |
| ENSDARG00000074283 | inpp5d | -2.44 | 4.71E-12 |
| ENSDARG00000069966 | alox5b.3 | 1.30 | 8.62E-12 |
| ENSDARG00000052826 | runx3 | -2.41 | 7.49E-11 |
| ENSDARG00000037861 | slc2a3b | -2.34 | 1.94E-10 |
| ENSDARG00000104820 | psd3l | -2.42 | 1.18E-09 |
| ENSDARG00000076534 | si:ch211-14a17.10 | -3.59 | 1.28E-08 |
| ENSDARG00000018263 | pdia2 | 2.73 | 1.05E-07 |
| ENSDARG00000057273 | alox5a | -2.37 | 3.96E-07 |
| ENSDARG00000077673 | nlrp16 | -6.32 | 9.60E-07 |
| ENSDARG00000073686 | heatr6 | 2.17 | 2.55E-06 |
| ENSDARG00000068745 | map4l | -2.40 | 8.18E-06 |
| ENSDARG00000092862 | si:ch211-260p9.3 | -2.35 | 9.11E-06 |
| ENSDARG00000022845 | lias | 3.25 | 9.80E-06 |
| ENSDARG00000067672 | card9 | -3.64 | 1.02E-05 |
| ENSDARG00000056258 | cdc27 | 1.24 | 3.74E-05 |
| ENSDARG00000094732 | mical3b | -2.85 | 3.88E-05 |
| ENSDARG00000045230 | cox6b1 | 1.12 | 7.34E-05 |
| ENSDARG00000100809 | g6fl | 1.58 | 8.46E-05 |
| ENSDARG00000059933 | plpp3 | 1.03 | 1.30E-04 |
| ENSDARG00000062956 | dagla | -2.59 | 2.12E-04 |
| ENSDARG00000059925 | usp24 | 1.01 | 2.12E-04 |
| ENSDARG00000089582 | si:dkey-265e15.2 | -2.93 | 2.31E-04 |
| ENSDARG00000013838 | sulf2b | -2.95 | 4.49E-04 |
| ENSDARG00000116660 | pigrl2.3 | -2.35 | 5.11E-04 |
| ENSDARG00000042641 | cyp51 | 1.48 | 5.66E-04 |
| ENSDARG00000063527 | elmo2 | -4.40 | 6.38E-04 |
| ENSDARG00000036776 | aldh8a1 | -2.75 | 9.79E-04 |
| ENSDARG00000102097 | nfkbiz | -2.42 | 1.13E-03 |

| | | | |
|---|---|---|---|
| ENSDARG00000044774 | pou5f3 | -2.77 | 1.44E-03 |
| ENSDARG00000018206 | nck2a | 2.55 | 1.56E-03 |
| ENSDARG00000036728 | si:dkey-211g8.1 | -4.17 | 2.02E-03 |
| ENSDARG00000021948 | tnc | -2.70 | 2.62E-03 |
| ENSDARG00000014386 | galnt6 | -2.47 | 2.68E-03 |
| ENSDARG00000104890 | si:ch211-76m11.3 | 6.07 | 5.99E-03 |
| ENSDARG00000090038 | BX248410.1 | -2.72 | 6.08E-03 |
| ENSDARG00000059832 | adgrb3 | -2.76 | 6.48E-03 |
| ENSDARG00000077710 | nlgn1 | -3.42 | 8.15E-03 |
| ENSDARG00000090164 | si:ch73-362m14.2 | -2.90 | 8.32E-03 |
| ENSDARG00000057378 | selenou1a | 1.25 | 9.60E-03 |
| ENSDARG00000060631 | GARNL3 | 3.66 | 1.18E-02 |
| ENSDARG00000062370 | bcl2l13 | 1.02 | 1.40E-02 |
| ENSDARG00000016470 | anxa5b | -2.52 | 1.49E-02 |
| ENSDARG00000099470 | muc5.3 | 1.57 | 1.70E-02 |
| ENSDARG00000003635 | mogat3b | -2.32 | 1.71E-02 |
| ENSDARG00000074590 | wdpcp | 2.36 | 1.86E-02 |
| ENSDARG00000102395 | CABZ01063602.1 | 3.48 | 2.07E-02 |
| ENSDARG00000095142 | si:ch211-208h16.4 | 2.63 | 2.27E-02 |
| ENSDARG00000004318 | cbwd | 1.30 | 2.31E-02 |
| ENSDARG00000044688 | dusp4 | 1.03 | 2.32E-02 |
| ENSDARG00000090369 | zgc:86896 | 1.12 | 3.17E-02 |
| ENSDARG00000079900 | shroom4 | 1.12 | 3.62E-02 |
| ENSDARG00000101214 | pkd1l2b | 1.21 | 3.81E-02 |
| ENSDARG00000068400 | znf131 | 1.72 | 4.40E-02 |
| ENSDARG00000079946 | sqlea | 1.09 | 4.56E-02 |
| ENSDARG00000058557 | il11b | 1.05 | 5.54E-02 |
| ENSDARG00000008278 | rcor2 | 1.47 | 6.54E-02 |
| ENSDARG00000096849 | si:dkey-16p21.8 | 1.13 | 6.64E-02 |

**Supplemental Table 5 – Statistics of the difference in expression of selected erythroid genes in cells treated with Epo, Dex, Kitlga compared to cells treated with Epo, Dex only.**

| Ensembl Accession ID | symbol | $log_2$FoldChange | p-adjusted |
|---|---|---|---|
| ENSDARG00000002194 | bcl2l13 | 1.02 | 1.40E-02 |
| ENSDARG00000003462 | rhd | 0.84 | 5.84E-03 |
| ENSDARG00000006818 | tfr1a | 0.55 | 1.06E-02 |
| ENSDARG00000008840 | hbae1.1 | 0.54 | 1.77E-01 |
| ENSDARG00000010252 | gata1a | 0.52 | 5.65E-02 |
| ENSDARG00000013477 | bcl2l10 | 0.52 | 1.18E-01 |
| ENSDARG00000017400 | epb41b | 0.49 | 3.78E-02 |
| ENSDARG00000019930 | hmbsa | 0.48 | 6.30E-02 |
| ENSDARG00000024295 | slc11a2 | 0.45 | 9.81E-02 |
| ENSDARG00000026766 | hbaa1 | 0.38 | 3.55E-01 |
| ENSDARG00000029019 | ppox | 0.38 | 1.89E-01 |
| ENSDARG00000030490 | klf1 | 0.37 | 2.68E-01 |
| ENSDARG00000062370 | sptb | 0.35 | 1.71E-01 |
| ENSDARG00000075641 | urod | 0.35 | 2.19E-01 |
| ENSDARG00000088330 | jak3 | 0.33 | 4.21E-01 |
| ENSDARG00000097011 | fech | 0.32 | 2.76E-01 |
| ENSDARG00000101322 | steap3 | 0.32 | 4.57E-01 |
| ENSDARG00000102167 | tal1 | 0.32 | 3.54E-01 |

**Supplemental Table 6. GO enrichment analysis in in cells treated with Epo, Dex, Kitlga compared to cells treated with Epo, Dex only.**

**Biological processes**

| Accession | GO.Term | Odds.Ratio | P.value |
|---|---|---|---|
| GO:0000028 | ribosomal small subunit assembly | 60 | 6.34E-12 |
| GO:0002181 | cytoplasmic translation | 53.8 | 2.63E-18 |
| GO:0000027 | ribosomal large subunit assembly | 50 | 1.38E-12 |
| GO:0042541 | hemoglobin biosynthetic process | 38.4 | 4.01E-07 |
| GO:0030218 | erythrocyte differentiation | 31.1 | 3.08E-15 |
| GO:0006412 | translation | 30.8 | 1.61E-73 |
| GO:0048821 | erythrocyte development | 22.7 | 9.68E-06 |
| GO:0043009 | chordate embryonic development | 18.7 | 1.55E-18 |
| GO:0006414 | translational elongation | 17.8 | 2.65E-05 |
| GO:0035162 | embryonic hemopoiesis | 15.1 | 2.91E-05 |
| GO:0051726 | regulation of cell cycle | 12 | 1.42E-06 |

**Cellular compartment**

| Accession | GO.Term | Odds.Ratio | P.value |
|---|---|---|---|
| GO:0022625 | cytosolic large ribosomal subunit | 77 | 1.22E-52 |
| GO:0022627 | cytosolic small ribosomal subunit | 72.9 | 9.55E-37 |
| GO:0015935 | small ribosomal subunit | 70 | 1.05E-10 |
| GO:0005840 | ribosome | 47.7 | 7.56E-82 |
| GO:0015934 | large ribosomal subunit | 41.6 | 3.67E-06 |

**Molecular function**

| Accession | GO.Term | Odds.Ratio | P.value |
|---|---|---|---|
| GO:0003735 | structural constituent of ribosome | 46.7 | 9.27E-80 |
| GO:0019843 | rRNA binding | 31.8 | 1.67E-07 |
| GO:0003723 | RNA binding | 5.14 | 5.75E-11 |
